# Symbiosis increases population size and buffers environmental fluctuations in a physiologically-structured model parameterized for thyasirid bivalves

**DOI:** 10.1101/2021.06.02.446784

**Authors:** Joany Mariño, Suzanne C. Dufour, Amy Hurford

## Abstract

Symbioses whereby one partner provisions a nutritional resource to the other may alter energy allocation towards reproduction and survival in the recipient partner, potentially impacting population dynamics. Asymbiotic thyasirid bivalves feed predominantly on free-living bacteria, which fluctuate in abundance due to seasonality-driven temperature variations. Symbiotic thyasirids are mixotrophs, gaining nutrients from free-living bacteria and symbiotic bacteria that they host on their enlarged gills. Symbiotic bacteria may function as an additional energy reserve for thyasirids, allowing the hosts to allocate more energy to reproduction. We hypothesize that, for symbiotic thyasirids, the symbionts are a nutritional source that mitigates resource limitation. Using Dynamic Energy Budget theory, we built a physiologically-structured population model assuming equal mortality rates in both species. We find that without seasonal fluctuations, symbiotic thyasirids have higher abundances than asymbiotic thyasirids since the symbionts increase reproduction. Both species have similar population sizes in fluctuating environments, suggesting different adaptations to seasonality: asymbiotic thyasirids have adapted their physiology, while symbiotic thyasirids have adapted through mixotrophy. Our results highlight the significance of linking individual energetics and life-history traits to population dynamics and are the first step towards understanding the role of symbioses in population and community dynamics.

## 1. Introduction

Nutritional symbiosis is a prevalent interaction that can increase the metabolic capabilities of the host [1, 2]. Hence, symbiosis has the potential to affect host life-history traits, such as fecundity and survival, which, in turn, determine population dynamics. However, how symbiosis can influence host ecology and how this would be translated into population and community dynamics is not known [3]. Given the ubiquity of nutritional symbiosis, disentangling how the effects arising from the lower trophic level (i.e., the symbionts) can determine the population structure of higher trophic levels (i.e., the hosts), in both constant and heterogeneous environments, is crucial to understanding the conditions that lead to the persistence of populations and communities [4].

Environmental heterogeneity and the pattern of environmental variation are thought to be determinant factors in the evolution of the niche breadth, particularly for traits such as foraging strategies [5, 6, 7]. Theory suggests that selection favours generalist strategies in populations experiencing environmental heterogeneity [6, 8, 9]. Experimental results have confirmed such findings [10, 11, 12]. For example, selection experiments in *Chlamydomonas* in constant environments have led to the evolution of specialists, either autotrophic or heterotrophic. Conversely, in temporally varying environments, selection favours generalists capable of both autotrophic and heterotrophic nutrition. However, in spatially varying environments, both specialists can be retained in the population [10, 11, 12]. Broadly, these results suggest that ecological specialists tend to be selected in environments that are homogeneous in space or time [9]. In contrast, a mix of strategies can coexist in spatially variable environments, while temporally varying environments tend to favour generalists [5, 6, 8, 13].

Nutritional symbioses in which the host has a mixotrophic nutrition (i.e. the host can combine the nutritional input from the symbionts with heterotrophic or autotrophic feeding) [14] can be considered to be generalist feeding strategies. For instance, mixotrophic symbioses are frequent in marine suspension feeders, which live in environments where light and plankton concentrations are variable and often limiting [15, 16]. In octocorals, such trophic flexibility has been proposed to maximize nutrient uptake, allowing for increased energy acquisition relative to asymbiotic species [17, 16]. Moreover, losing the symbionts may not significantly affect the energetic input of a mixotrophic individual, making it less affected by environmental variability [14, 18, 19, 20, 21]. Thus, in seasonal environments, symbionts can provide energy to a mixotrophic host and stabilize the discontinuous energy inputs in resource availability [14, 17, 18].

Another notable example of mixotrophy occurs in symbiotic thyasirid bivalves, a family that stands out for including symbiotic as well as asymbiotic members [22, 23]. Thyasirid bivalves, known as cleft or hatchet clams, have a cosmopolitan distribution in marine waters, living infaunally in the deep sea (i.e. cold seeps, whale falls and hydrothermal vents) or in soft, oxygen-poor and sulphide-rich sediments (e.g., sewage enriched sediments, fjords) [1, 24, 25]. Thyasirids are most abundant in sediments that have a sparse infauna, which has been attributed to the bioturbation that other burrowers produce, that disrupts the tunnel system and decreases the ability of the thyasirids to obtain sulphide [26, 27]. Thyasirids exhibit a variety of feeding strategies. Some asymbiotic species mainly rely on suspension feeding, obtaining nutrients through an inhalant tube constructed with their foot [28, 29]. Other asymbiotic thyasirids are deposit feeders, using the surface of the foot first to collect particulate organic matter, transfer it into the mantle cavity and then deposit it in the mouth [30]. Symbiotic thyasirids are flexible mixotrophs: they have nutritional input from suspension feeding but mainly rely on combining deposit feeding on free-living bacteria with predation on their bacterial symbionts, which are periodically endocytosed and digested [31]. Symbiotic thyasirids appear to consume their symbionts depending on environmental conditions, particularly the presence of sulphide and particulate food in the sediment [32, 33]. Symbiotic thyasirids are flexible mixotrophs that can digest their symbionts depending on environmental conditions, particularly the presence of sulfide and external particulate food [32, 33]. Evidence in other bivalves suggests that changes in the relative importance of different food sources are likely correlated to particulate food abundance [34]. A mixotrophic nutrition is considered a strategy that allows symbiotic thyasirids to thrive in fluctuating environments [33, 35].

In the fjord of Bonne Bay (Newfoundland, Canada), two small species of thyasirids (∼3.5 mm in length) are sympatric and have a patchy distribution. The first species resembles the northern hatchet-shell, *Thyasira gouldi* (in shell char-acteristics and internal anatomy), and therefore is referred to as *T*. cf. *gouldi*; the second species, *Parathyasira* sp., is asymbiotic. Both symbiotic *T*. cf. *gouldi* and asymbiotic *Parathyasira* are particulate feeders that rely on chemoautotrophic bacteria as their primary resource (60% and 70%, respectively), with lesser contributions of suspended and particulate organic matter [29]. However, rather than collecting chemoautotrophic bacteria from sediments through pedal feeding, *T*. cf. *gouldi* harbours these bacteria extracellularly as symbionts on enlarged gills and digests them as an additional resource [29, 25]; hence, it is considered a mixotrophic species. For *Thyasira sarsi*, another mixotrophic thyasirid, between 26% and 76% of their nutrition has been estimated to be obtained from the bacterial symbionts [32]. Given that the carbon isotope composition in *T*. cf. *gouldi* overlaps the lower range of the isotopic signature of *T. sarsi* [32, 29], it is likely that the reliance of *T*. cf. *gouldi* on symbionts may be similar to the lower limit of *T. sarsi*’s, comprising approximately 25% of their diet. For thyasirids inhabiting an environment with strong seasonality, temperature and resource fluctuations will affect the individual metabolic rates and the costs associated with maintaining the symbionts. Previous theoretical research showed that symbiotic *T*. cf. *gouldi* has a smaller energy reserve, which implies reduced energy assimilation and mobilization fluxes, lower somatic maintenance costs and growth rate, and more significant energy allocation to maturity and reproduction [36]. However, how the nutritional differences between symbiotic *T*. cf. *gouldi* and asymbiotic *Parathyasira* are reflected at the population and community levels is not known.

Some previous results support the hypothesis that a mixotrophic (generalist) strategy results in higher energy allocation to reproduction [18, 37, 38, 39]. Thus, in a constant environment, a mixotrophic population should have larger abundances than the asymbiotic (specialist) population. Here, we hypothesize that when there is seasonality, relying on symbionts will buffer the fluctuations in resource availability for the host population. We predict that if symbionts effectively mitigate resource variability, then the mixotrophic population will be less prone to extinction during winter when the abundance of free-living bacteria becomes limiting. Our prediction should hold while the abundance of symbionts is not zero and is at least equal to the lowest free-living bacterial abundance. However, the buffering effect of the symbionts should decrease as the host becomes more specialized and increases reliance on the symbionts. Hence, in highly seasonal environments, a low or intermediate level of dependence on symbionts (i.e. a generalist strategy) should be favoured over the specialist strategy.

Since the physiological responses of individuals can be considered the underlying basis of their ecological dynamics, models that consider the organismal bioenergetics are powerful tools to understand how energetic constraints determine changes in the niche of a species as a consequence of environment fluctuations [40]. Dynamic Energy Budget (DEB) models describe the rates at which an individual assimilates and uses energy for maintenance, growth, and reproduction [43, 44]. DEB theory provides a characterization of the life cycle of an organism through a model that describes the links between the metabolic processes through-out the lifespan of the individual [40]. Among its many applications, DEB theory has been proposed as a natural approach to describe systems that involve internal symbionts [41], or those constrained by environmental fluctuations [42].

Previously, we suggested that the energy allocation patterns in symbiotic thya-sirids may represent an evolutionary strategy where the symbionts function as a partial energy reserve, allowing the individuals to invest more energy in reproduction [36]. To understand how the differences due to feeding and symbiosis translate to the population level and shape host population dynamics, we combine the individual-level energy budget dynamics with a physiologically-structured population model [44, 45]. We built a population model that accounts for the species’ physiology according to the individual DEB model and takes into account the seasonal pattern of temperature and resource abundance. Using this model, we simulated the dynamics of the symbiotic *T*. cf. *gouldi* and the asymbiotic *Parathyasira*. We focus on the differences in populations inhabiting a constant versus seasonal environment, and evaluate different scenarios of symbiont dependence and abun-dance. We show how the symbiotic strategy is likely to mitigate the effects of environmental variability in a population of symbiotic thyasirids. We discuss the buffering effect of the symbionts in terms of the evolution and ecological adaptation of thyasirids and mixotrophic bivalves.

## 2. Materials and methods

### 2.1. Overview

We formulated a continuous population model that focuses on the representation of individual physiology and life history. To describe individuals, we used the DEB-abj model [46], which includes the larval stage of bivalve molluscs and is structured by energy, volume, maturity and reproduction (Fig. 1). We used published data from *T*. cf. *gouldi* and *Parathyasira* sp. to parameterize the model [36]. We assumed that individuals in the population could exploit one or two resources, depending on whether they are asymbiotic or symbiotic. To explicitly include the dependency of the resource on the environmental temperature, we modelled the resource according to relationships derived from the Metabolic Theory of Ecology [47]. To test our predictions of how symbiosis affects populations of thyasirid bivalves, we conducted numerical simulations for *T*. cf. *gouldi* and *Parathyasira* in different environmental conditions that consider various temperature and resource availability scenarios. To further analyze the possible effects of symbiosis on populations of *T*. cf. *gouldi*, we carried out simulations representing different relative symbiont abundances and distinct contributions of symbionts to the host’s diet.

**Figure 1.**
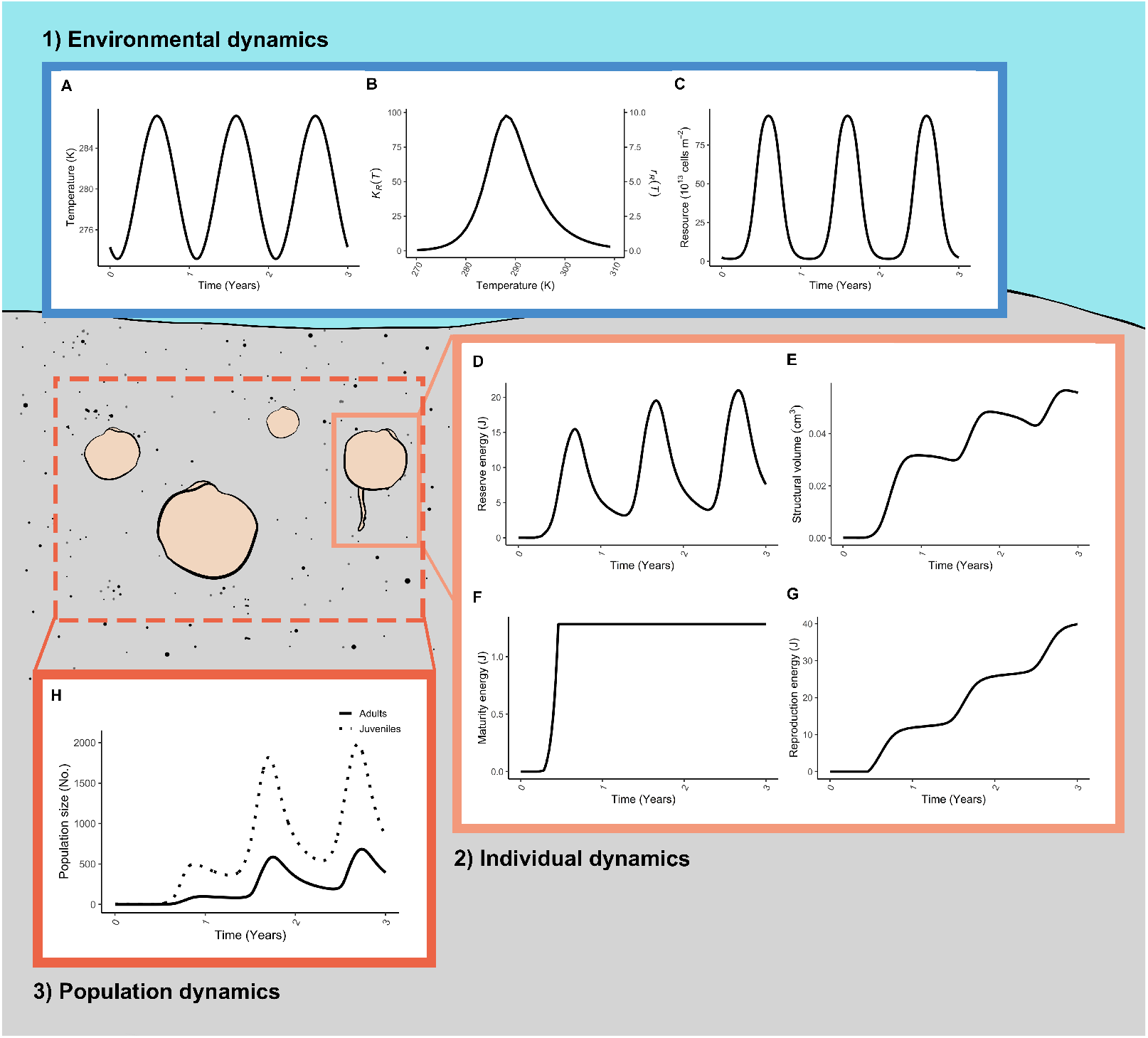
Representation of the three main components of the model for *Thyasira* cf. *gouldi* in time: 1) the environment (A, B, C), 2) the individual (D, E, F, G) and 3) the population (H). The environmental variables of temperature (A, equation 1) and free-living bacterial resource (C, equation 2) determine each individual’s dynamics. The carrying capacity of the free-living bacte-rial resource (*K*_*R*_(*T*)) and its growth rate (*r*_*R*_(*T*)) are functions of the environmental temperature (B, equations 3 and 4). For both parameters, we follow relationships derived from the Metabolic Theory of Ecology [47]. We characterized the individuals by four variables: energy in reserve (D), structural volume (E), maturity (F) and reproduction energy (G), according to the DEB model (equation 7). We obtained the population dynamics (H) by numerically integrating over all the individuals (equation 10). The simulations’ parameters are given in Tables 2 and 3. We assumed an initial condition of 5 embryos for the simulation of the population.

In the next sections, we present first our description of the environment (section 2.2), consisting of the temperature and food resource. Next, we explain the individual dynamics at the core of the DEB modelling approach (section 2.3), considering feeding, growth and reproduction. Using these components, we then define the population dynamics (section 2.4) and describe the implementation of the model (section 2.5).

### 2.2. Environment

#### 2.2.1. Temperature

We modelled an annual cycle that corresponds to the seafloor temperatures at Bonne Bay, which range from 0.7 to 14 °C, approximately [25, see Fig. 1A]. More specifically, the temperature at time *t* oscillates around the average temperature (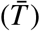) with amplitude *T*_*a*_ and period equal to the length of the year:

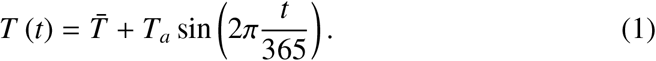

For comparative purposes, we also modelled a constant environment assuming that the temperature is equal to the mean annual temperature, 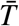.

#### 2.2.2. Resource

The primary resource for *T*. cf. *gouldi* and *Parathyasira* is free-living, chemoautotrophic bacteria [29]. In cold, marine sediments, such as Bonne Bay, bacterial production is typically seasonal, with specific growth rates increasing with temperature; further, bacterial production is directly proportional to bacterial biomass [48]. Thus, we assumed that the free-living bacterial resource *R* is a function of the environmental temperature *T*, and follows logistic growth:

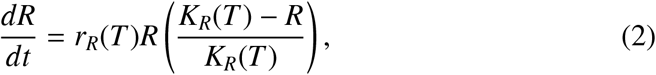

where *r*_*R*_ is the resource growth rate and *K*_*R*_ is the maximum resource density. For notational simplicity, we write *T* instead of *T* (*t*); however, it should be understood that temperature may be a function of time as described in equation 1.

To include the dependence of the resource on the environmental temperature, we described both resource parameters using relationships derived from the Metabolic Theory of Ecology [47]. We assumed that the growth rate and carrying capacity increase exponentially with temperature. Further, around the limits of the bacteria’s thermal niche, both parameters drop steeply to zero, according to a Sharpe-Schoolfield term (see Fig. 1B) [49]. Hence, the resource growth rate is given by:

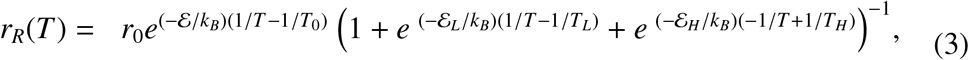

where *r*_0_ is the resource growth rate at a reference temperature *T*_0_, *k*_*B*_ is the Boltz-mann constant, *ε* is the average activation energy driving resource growth at intermediate temperatures, *ε* _*L*_ and *ε* _*H*_ are the inactivation energies that determine the slope of the resource growth rate as it drops to zero at the lower and upper thermal tolerance limits, *T*_*L*_ and *T*_*H*_, respectively.

The maximum resource density follows a similar formulation:

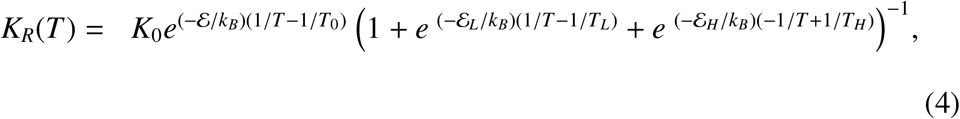

where *K*_0_ is the maximum resource density at a reference temperature *T*_0_. As with the resource growth rate, the parameters *ε, ε* _*L*_ and *ε* _*H*_ represent the temperature sensitivity of the maximum resource density within and outside of the lower and upper temperature thresholds, *T*_*L*_ and *T*_*H*_.

### 2.3. Individual dynamics

We described the individual life history and physiology (i.e. feeding, growth, and reproduction) as a function of the individual state variables and the state of the environment.

#### 2.3.1. Feeding

All individuals forage on the resource *R* (free-living bacteria) following a function *f* (*t, T, R*), which depends on the time of the year, the temperature and the resource abundance. We normalized the function according to the maximum resource *R*_*max*_, which always takes the maximum value at *T*_0_. Asymbiotic individuals feed only on one resource, such that the normalized feeding function is given by:

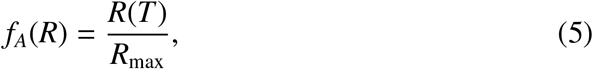

Symbiotic individuals can forage on the symbionts as an additional resource, which we modelled as a constant abundance *S* scaled by the free-living resource.

Thus, the normalized feeding function for symbiotic individuals *f*_*S*_ (*t, T, R*) includes their reliance on symbionts α, and is given by:

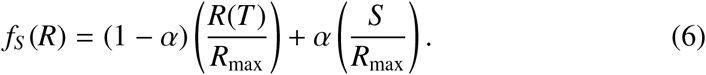

Here, α = 0 results in *f*_*A*_ and corresponds to asymbiotic feeding, while α = 0 represents an obligate feeding symbiosis.

#### 2.3.2. Growth and reproduction: DEB-abj model

The individual dynamics are according to the DEB theory [44]. This approach distinguishes between the biomass of the organism that functions as an energy reserve and as structure. Specifically, we use the DEB-abj model, which is a one parameter extension of the standard DEB model that accounts for a growth pattern recognized in bivalves termed metabolic acceleration [46].

Each individual is characterized by four state variables: energy in reserve (*E*), structural volume (*L*^3^), cumulative energy invested into maturation (*E*_*H*_), and cumulative energy invested into reproduction (*E*_*R*_). The dynamic of the individual in time is given by:

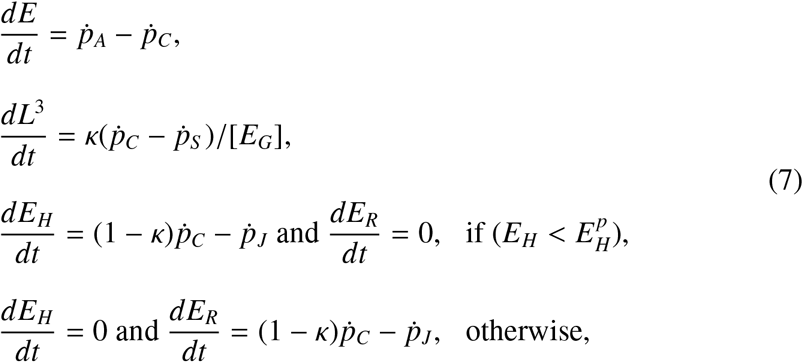

where the energy fluxes are denoted by 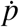 (see Table 1), κ is the fraction of mobilized reserve allocated to somatic metabolism, [*E*_*G*_] is the specific cost to grow one unit of structure, and 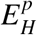 is the puberty threshold.

**Table 1.** Energy fluxes (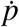,J/d) and shape correction function (ℳ) at each developmental stage. Each stage is defined according to the cumulative maturity thresholds 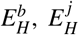 and 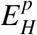, represent birth, metamorphosis, and puberty, respectively. The scaled functional response is *f* (0 ≤ *f* ≤ 1, where 1 is the highest amount of food), 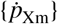 is the maximum surface-area specific ingestion rate 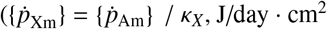, for a maximum surface-area specific assimilation rate 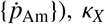 is the assimilation efficiency from food to reserve, 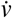 is the energy conductance rate from the energy reserve (cm/day), 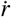 is the individual growth rate (1/day, equation 8), 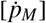is the volume-specific somatic maintenance cost (J/day · cm^3^), 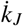is the maturity maintenance rate coefficient. The structural lengths at the beginning and at the end of the acceleration are *L*_*b*_ and *L*_*j*_, which correspond to the structural lengths at birth and at metamorphosis. Notation: square brackets ([ ]) indicate quantities related to structural volume, curly brackets ({ }) denote quantities related to structural surface-area, dots (·) indicate rates.

| Flux | Embryo<br>($E_H \leq E_H^b$) | Early juvenile<br>($E_H^b < E_H \leq E_H^j$) | Late juvenile<br>($E_H^j < E_H < E_H^p$) | Adult<br>($E_H \geq E_H^p$) |
| --- | --- | --- | --- | --- |
| Feeding, $\dot{p}_X$ | 0 | $f\{\dot{p}_{Xm}\}\mathcal{M}L^2$ | $f\{\dot{p}_{Xm}\}\mathcal{M}L^2$ | $f\{\dot{p}_{Xm}\}\mathcal{M}L^2$ |
| Assimilation, $\dot{p}_A$ | $\kappa_X \dot{p}_X$ | $\kappa_X \dot{p}_X$ | $\kappa_X \dot{p}_X$ | $\kappa_X \dot{p}_X$ |
| Mobilization, $\dot{p}_C$ | $E\dot{v}(\mathcal{M}/L - \dot{r})$ | $E\dot{v}(\mathcal{M}/L - \dot{r})$ | $E\dot{v}(\mathcal{M}/L - \dot{r})$ | $E\dot{v}(\mathcal{M}/L - \dot{r})$ |
| Soma maint., $\dot{p}_S$ | $[\dot{p}_M]L^3$ | $[\dot{p}_M]L^3$ | $[\dot{p}_M]L^3$ | $[\dot{p}_M]L^3$ |
| Maturity maint., $\dot{p}_J$ | $\dot{k}_J E_H$ | $\dot{k}_J E_H$ | $\dot{k}_J E_H$ | $\dot{k}_J E_H^p$ |
| Shape function, $\mathcal{M}$ | $L_b/L_b = 1$ | $L/L_b$ | $L_j/L_b$ | $L_j/L_b$ |

The growth rate for each individual is given by:

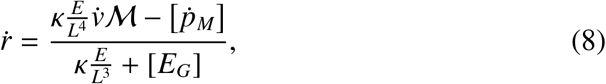

Where ℳ is a shape correction function that varies according to the stage of the individual (see Table 1), 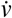 is the energy conductance rate, and 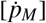 is the somatic maintenance cost.

The energy fluxes for all the metabolic rates are temperature-dependent (see Section 1.3 in [44]), therefore the parameters of the model are standardized to a reference temperature of 20°C. The correction between the reference temperature and the empirical temperature *T* is done through the Arrhenius relationship:

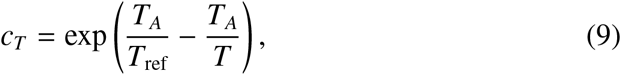

where *c*(*T*) is the correction factor for a certain temperature *T, T*_*A*_ is the Arrhenius temperature and *T*_*re f*_ is the reference temperature. For example, the mobilization flux (Table 1) at temperature *T* becomes: 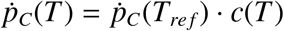.

### 2.4. Population dynamics

We represent the number of individuals by *n* ΔΩ, where ΔΩ is an interval of the four individual DEB (or i-state) variables (*n*(*t, E, V, E*_*H*_, *E*_*R*_)) and its dynamic in time is given by:

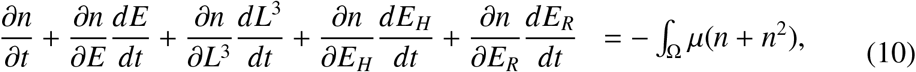

with a per capita mortality rate μ. The set of all the possible i-states defines the population state space Ω ⊂ ℝ^4^. To prevent individuals from leaving the domain, we included no-flux boundary conditions for each i-state variable: 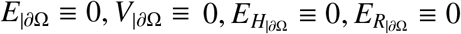.

The population state space is divided into subsets that represent the different life stages of the individuals. For simplicity, we group the two juvenile stages together, and consider the domains Ω_*J*_, and Ω_*A*_, corresponding to the juvenile and adult stages. The boundary between these subsets is given by the cumulative maturity energy threshold parameter 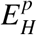. For biological realism, we set a no-flux boundary condition at 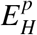 when 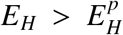. This ensures that once individuals reach the adult state, they cannot later revert to the juvenile state.

The reproduction of the adult population gives the boundary condition at age zero:

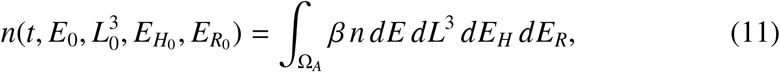

where we consider that the per capita survival rate of embryos β is a constant. Additionally, we assume that individuals are born in the population at the origin of the domain.

### 2.5. Model analysis

We implemented our population model in the R language [50] and solved it as an initial value problem through the methods of lines. For this, we discretized the evolution equations with finite differences and performed the time integration using the lsoda initial value problem solver from the package **deSolve** [51]. Our simulations represented experimental populations that start with 5 embryos, for either species (as in [52, 53]). Thus, the initial conditions for our numerical simulations were:

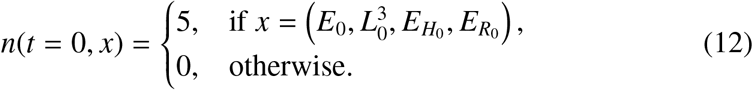

Using this implementation, we conducted three sets of simulations. First, to investigate the differences in the population dynamics between both species (section 3.1), we conducted simulations at constant and fluctuating environments. We assumed here that the abundance of the symbionts was equal to the mean abundance of the free-living bacterial resource 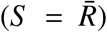. Further, we assumed that symbionts provide 25% of the diet of symbiotic individuals (α = 0.25), similar to *T. sarsi* [32]. Second, to analyse the effect of the abundance of symbionts in a host population inhabiting a fluctuating environment (section 3.2), we investigated three possible cases: i) the abundance of the symbionts is equal to the average abundance of the resource 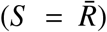 ; ii) the symbionts are more abundant than the resource (*S* = *R*_max_); and iii) the symbionts are less abundant than the resource (*S* = *R*_min_). Here, we also assumed that the contribution of symbionts to a host individuals’ diet is 25% (α = 0.25). Third, to assess how the reliance on symbionts alters the abundance of the host population (section 3.3), we evaluated the impact of the dependence on symbionts (α) for the case 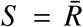. This way, we consider several values of α, ranging from individuals that do not rely on symbionts (α = 0), to those that rely solely on symbionts as their resource (α = 1).

The parameters that we used to conduct the simulations are given in Table 2, for the individual dynamics, and Table 3, for the environment and population. For both species, we estimated the parameters using allometric and life-history data, see [36] for details on the data and model parametrization. The results presented for these sections correspond to the values of the population after transient dynamics have disappeared.

**Table 2.** Parameter values for the individual-level dynamics (equation 7) for the symbiotic *Thyasira* cf. *gouldi* and the asymbiotic *Parathyasira* sp. [36]. Notation: square brackets ([ ]) indicate quantities related to structural volume, curly brackets (·) denote quantities related to structural surface-area, dots ({ }) indicate rates.

| Description | Symbol | *Thyasira* cf. *gouldi* | *Parathyasira* | Unit |
| --- | --- | --- | --- | --- |
| Maximum assimilation rate | $\{\dot{p}_{Am}\}$ | 1.427 | 2.547 | J/day · cm<sup>2</sup> |
| Assimilation efficiency | $\kappa_X$ | 0.8 | 0.8 | - |
| Energy conductance rate | $\dot{v}$ | 0.02 | 0.02 | cm/day |
| Allocation fraction to soma | $\kappa$ | 0.883 | 0.958 | - |
| Somatic maintenance cost | $[\dot{p}_M]$ | 15.78 | 23.61 | J/day · cm<sup>3</sup> |
| Maturity maintenance coefficient | $\dot{k}_J$ | 0.002 | 0.002 | 1/day |
| Specific cost for structure | $[E_G]$ | 2355 | 2348 | J/cm<sup>3</sup> |
| Maturity at birth | $E_H^b$ | 2.639e-4 | 7.193e-5 | J |
| Maturity at metamorphosis | $E_H^J$ | 0.011 | 0.002 | J |
| Maturity at puberty | $E_H^p$ | 1.283 | 0.96 | J |
| Weibull aging acceleration | $\dot{h}_a$ | 9.844e-8 | 1.262e-7 | 1/day<sup>2</sup> |
| Shape coefficient | $\delta_M$ | 0.507 | 0.64 | - |
| Arrhenius temperature | $T_A$ | 8000 | 8000 | K |
| Reference temperature | $T_{ref}$ | 293.15 | 293.15 | K |

**Table 3.** Parameter values for the environment and population-level dynamics for the symbiotic *Thyasira* cf. *gouldi* and asymbiotic *Parathyasira* sp. We assume that the parameter values are equal for both populations.

| Description | Symbol | Value | Unit | Reference |
| --- | --- | --- | --- | --- |
| *Environment* |  |  |  |  |
| Temperature | $T$ | variable | K | [25] |
| Mean temperature | $\bar{T}$ | 279.15 | K | [25] |
| Temperature amplitude | $T_a$ | 7 | - | [25] |
| Lower temperature limit for survival | $T_L$ | 273.15 | K | - |
| Upper temperature limit for survival | $T_H$ | 303.15 | K | - |
| Reference temperature | $T_0$ | 288.15 | K | $(T_H + T_L)/2$ |
| Mean activation energy | $\mathcal{E}$ | 0.43 | J | [47] |
| Lower tolerance limit inactivation energy | $\mathcal{E}_L$ | 1.9 | - | - |
| Upper tolerance limit inactivation energy | $\mathcal{E}_H$ | 1.9 | - | - |
| Boltzman constant | $k_B$ | $8.617e^{-5}$ | eV/K | - |
| Free-living bacterial resource | $R$ | variable | - | - |
| Free-living bacterial resource growth rate at $T_0$ | $r_0$ | $5e^6$ | day<sup>-1</sup> | - |
| Free-living bacterial resource density at $T_0$ | $K_0$ | $5e^7$ | day<sup>-1</sup> | - |
| Maximum free-living bacterial resource | $R_{max}$ | variable | - | - |
| Symbiont scaled abundance | $S$ | variable | - | - |
| *Population* |  |  |  |  |
| Survival rate of embryos | $\beta$ | 0.005 | day<sup>-1</sup> | [54] |
| Death rate | $\mu$ | 0.001 | day<sup>-1</sup> | [54] |

## 3. Results

### 3.1. The symbiotic population has a greater proportion of adults

In the constant environment, both the asymbiotic and the symbiotic (α = 25%) species reach carrying capacity and stabilize after two years (Fig. 2A). Both populations are dominated by individuals in the juvenile classes, with all the stages following the same growth pattern (Fig. 2B, C). However, the symbiotic *T*. cf. *gouldi* population shows a faster growth rate and reaches a higher carrying capacity, relative to the asymbiotic *Parathyasira* (Fig. 2A-C) since individuals of *T*. cf. *gouldi* allocate more energy to reproduction [36]. Further, in the symbiotic population, a larger proportion of individuals are in the adult stage (Fig. 2C).

**Figure 2.**
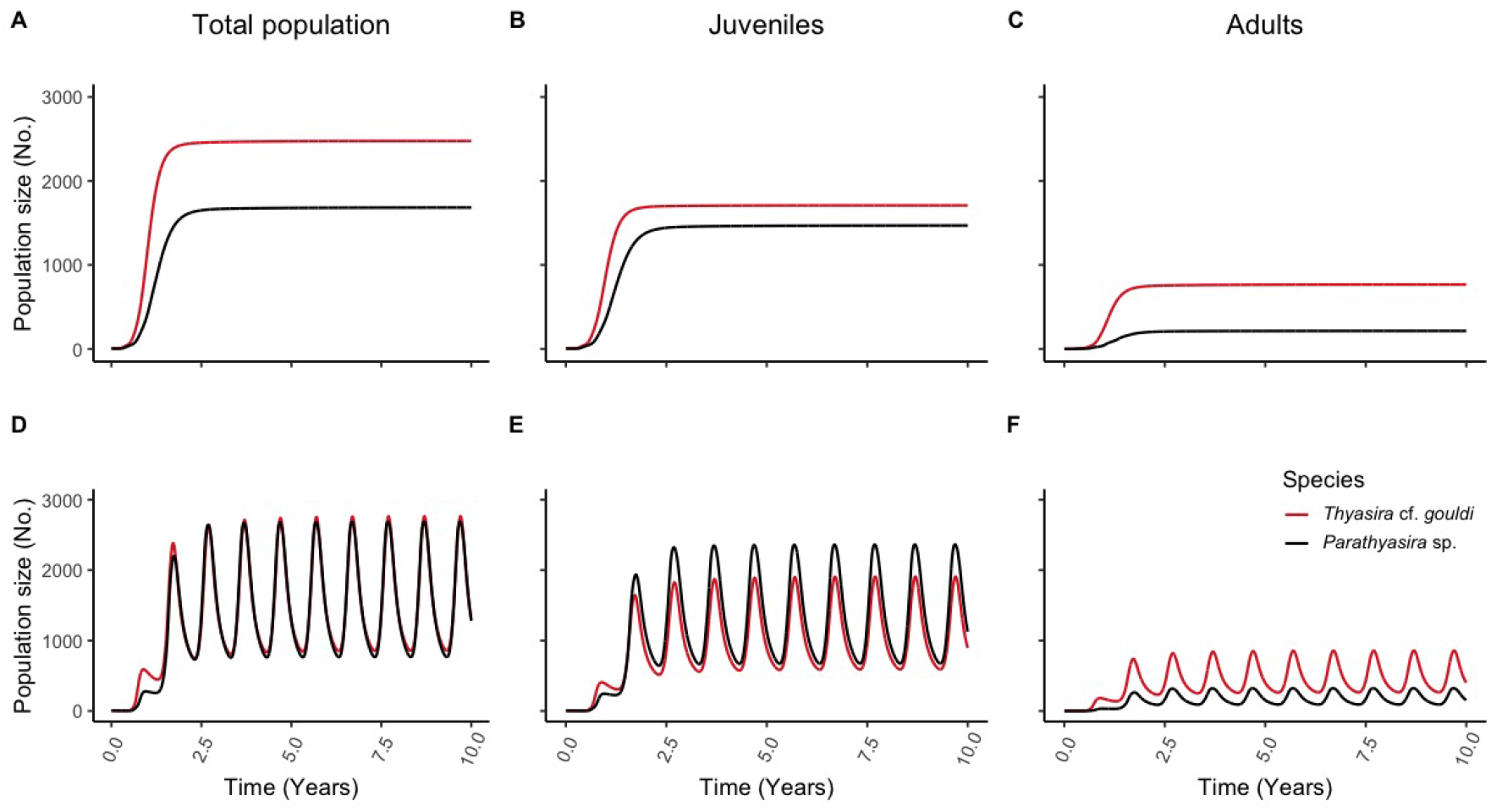
Modelled long term population dynamics for the symbiotic *T*. cf. *gouldi* and asymbiotic *Parathyasira* sp. in a constant (A-C) or fluctuating environment (D-F). For all cases, the initial population consists of five embryos. The reliance of *T*. cf. *gouldi* on symbionts is assumed to correspond to 25% of their diet. The symbiotic strategy allows individuals to invest more energy in reproduction, resulting in larger population sizes than the asymbiotic population in the constant environment (A). In the seasonal environment (D), both populations exhibit yearly cycles due to the combined effect of temperature and resource fluctuations. The stage dynamics for juvenile individuals (B, E) and adult individuals (C, F) exhibit the same pattern as the total population, with juveniles dominating both species’ populations. In both environment scenarios, the proportion of adults is greater in the symbiotic population.

In the seasonal environment, the populations of both species experience yearly cycles of low temperatures and low abundances of free-living bacteria (Fig. 1A, B), which cause a decrease in the individual growth and maturity rates as well as in the production of offspring (Fig. 1D, F). Consequently, both populations exhibit similar amplitude fluctuations with a one-year periodicity (Fig. 2D). During the periods of low free-living bacteria abundance, the symbiotic and asymbiotic thyasirid populations have similar sizes (Fig. 2D). Individuals in all the stages follow the same regular oscillations, with juveniles being the most abundant class in both populations (Fig. 2E). However, the proportion of adults is higher in the symbiotic population, reaching a greater abundance and a larger minimum size than the asymbiotic population (Fig. 2F).

### 3.2. Increasing symbiont abundance increases the population size of the host

In the seasonal environment, the dynamics of the population of symbiotic *T*. cf. *gouldi* vary according to the abundance of symbiotic bacteria (Fig. 3A). The population attains the largest size when the abundance of symbionts is greater than the abundance of the free-living bacteria (i.e. *S* > *R*, Fig. 3A). In this case, the mean population size and the amplitude of the yearly cycles are larger than in the other scenarios. As the abundance of the symbionts decreases (i.e. *S* < *R*), the host population exhibits cycles of smaller amplitudes and reaches a smaller mean size (Fig. 3A).

**Figure 3.**
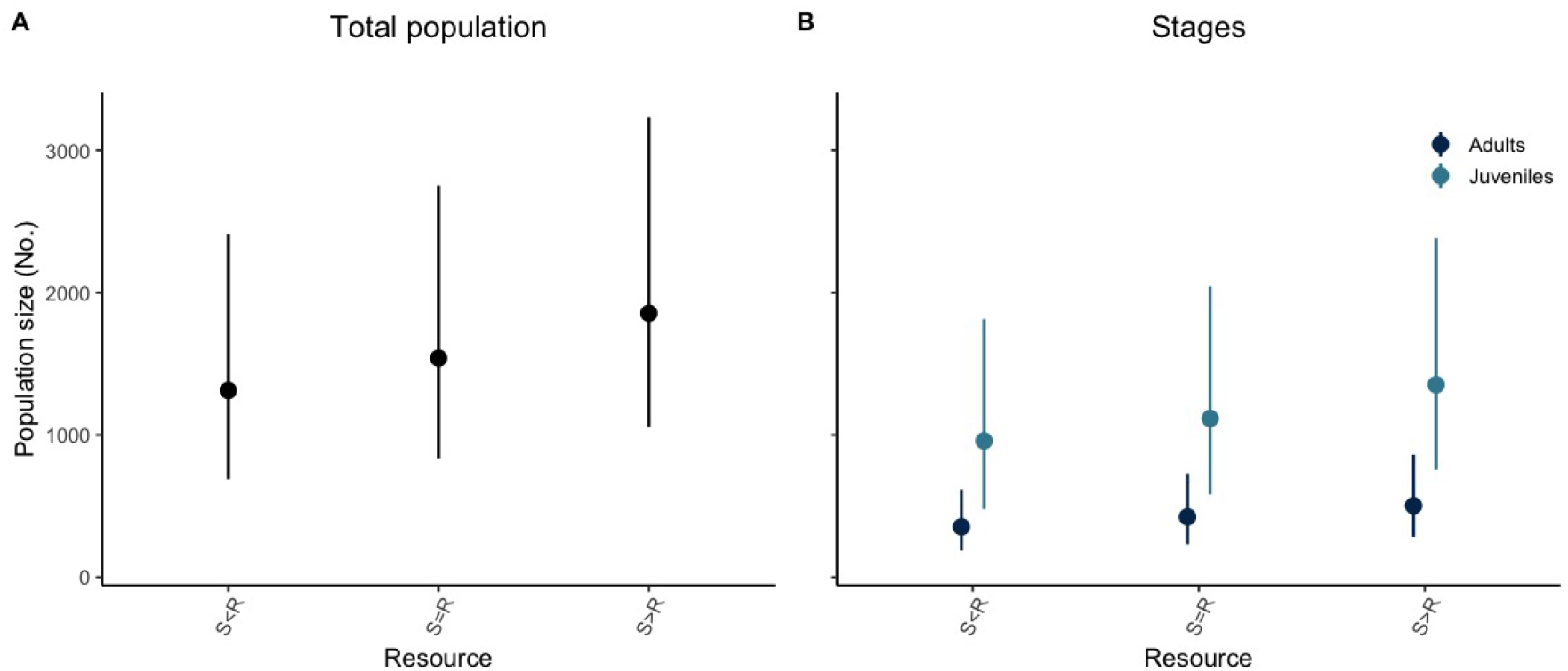
Effect of different symbiont abundances (*S*) relative to the free-living bacteria (*R*) on the host population of *T*. cf. *gouldi* in a seasonal environment. For all cases, the initial population consists of five embryos, and the reliance of *T*. cf. *gouldi* on symbionts corresponds to 25% of their diet. Points represent the average population size after discarding transient dynamics, and lines correspond to the yearly cycle’s amplitude. An increasing relative symbiont abundance is predicted to increase the average, minimum and maximum population sizes in *T*. cf. *gouldi* (A). The stage dynamics exhibit the same pattern as the total population, with juveniles dominating both species’ populations (B).

In the three scenarios of symbiont abundance that we considered, the dynamics of the different age classes of symbiotic *T*. cf. *gouldi* follow the same pattern as the population (Fig. 3B). Individuals in the juvenile stage dominate the populations and exhibit the largest amplitude in abundance when the conditions for the symbionts are favourable. In contrast, fluctuations in amplitude are smaller for the adult stage. Nevertheless, an increase in the abundance of symbionts favours a higher mean number of adults.

### 3.3. Increasing reliance on symbionts reduces the amplitude of the population cycles of the host

For populations of symbiotic *T*. cf. *gouldi* in a seasonal environment, the amplitude of the yearly cycles depends on the reliance of each individual host on the symbiotic bacteria (Fig. 4A). The largest mean population size and annual cycles with the highest amplitude occur in the population where individuals do not obtain nutrients from symbionts (i.e. α = 0, equivalent to asymbiotic individuals, Fig. 4A). An increasing reliance on symbionts reduces the effect of seasonal fluctuations in the free-living bacterial resource. Consequently, both the amplitude of the population cycles and the mean population size decrease. The smallest amplitudes and mean population sizes occur in the population where hosts rely entirely on symbionts (i.e. α = 1, equivalent to obligate symbiosis), given that they are not affected by fluctuations in resource availability. However, populations that rely entirely on symbionts still experience fluctuations in abundance since temperature affects the metabolic rates. Furthermore, the minimum population size rises with increasing reliance on symbionts, the highest minimum occurring in the population where individuals are obligate symbionts. Therefore, in a seasonal environment, populations with a greater dependence on symbionts are less likely to experience extinction.

**Figure 4.**
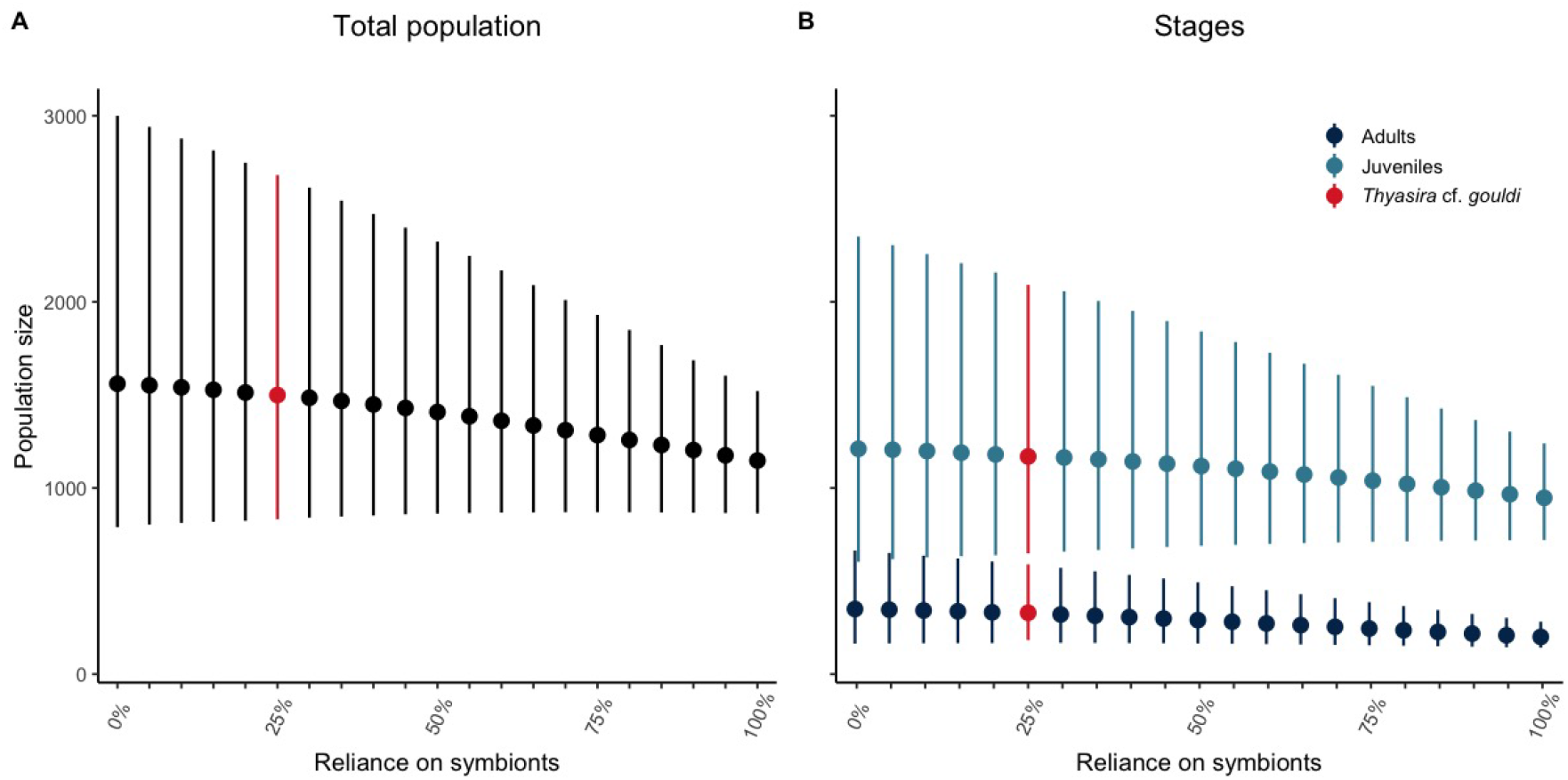
Effect of varying symbiont reliance (α) on the host populations in a seasonal environment. For all simulations, the initial population consists of five embryos. *T*. cf. *gouldi*, which has a dependence on symbionts assumed to be 25%, is highlighted for reference. Points represent the average population size after discarding transient dynamics, and lines correspond to the amplitude of the yearly cycle. An increasing symbiont dependence decreases the magnitude of the annual population cycles (A). As dependence on symbionts increases, the average and maximum population sizes decrease; however, the minimum population size increases. The stage dynamics exhibit the same pattern as the total population, with juveniles dominating both species’ populations (B).

As in our previous results, the dynamics of the two stages of symbiotic *T*. cf. *gouldi* follow the same fluctuating pattern as the combined population (Fig. 4B). In all the scenarios considered, juvenile individuals predominate and show the greatest variation in abundance, whereas individuals in the adult class exhibit cycles of smaller amplitude. Both the adult and juvenile stages reach a greater average population size when they have a low reliance on symbionts. Similarly, the mean population size of both stages decreases with an increase in the individual specialization on the symbionts. For the two stages, the minimum population size shows an increase as the dependence on symbionts increases.

## 4. Discussion

We show how the host’s individual physiology and the abundance of and dependence on symbionts affect thyasirid population dynamics in constant and seasonal environments. Our simulations for a constant environment suggest that the mixotrophic species may reach a higher population size than the asymbiotic species. In a seasonal environment, the population of symbiotic adults has a higher growth rate during periods of the year with higher temperatures and a greater abundance of free-living resources. Moreover, the symbiont abundance and the degree of specialization in the host’s diet modulate the effect of symbiosis. Thus, our results support our initial hypothesis that symbiosis with an intermediate level of reliance on the symbionts mitigates the impact of resource seasonality.

### 4.1. Population dynamics

We found that symbiotic thyasirids reach larger population sizes and have faster population growth than the asymbiotic species in a constant environment (Fig. 2A). A similar pattern has been found in *T. sarsi*, which has an intermediate reliance on bacterial symbionts (between 26% to 76%) and exhibits a faster population growth rate, reaching larger population sizes when compared to the sympatric and less symbiont dependent *T. equalis* (which has a reliance below 26%) [32]. However, unlike *T. equalis*, the asymbiotic *Parathyasira* have higher somatic growth rates and reach larger sizes at maturity, relative to the symbiotic *T*. cf. *gouldi* [36]. Further, our results show that both populations are composed mostly of juvenile individuals (Fig. 2B, E). A comparable population structure has been observed in *Thyasira gouldi*, which exhibits a bimodal distribution year-round [55]. Our findings for a seasonal environment show that both populations experience yearly cycles. Similarly, for *Thyasira gouldi, T. sarsi*, and *T. equalis* empirical data has suggested that they have variable population sizes [55, 24]. Thus, despite the limited empirical evidence from other thyasirid bivalves or symbiotic animals, our results broadly agree with the literature.

### 4.2. Effect of symbiosis on the host population

In our simulations, the symbiotic population of *T*. cf. *gouldi* experiences os-cillations of a similar amplitude relative to the asymbiotic population (Fig. 2D). When we consider a symbiotic population in which the symbionts are more abun-dant, the amplitude of the population cycles increases (Fig. 3). Conversely, when the individuals have a higher dependency on the symbionts, the yearly cycles have smaller amplitudes (Fig. 4). Broadly, our results indicate that the magnitude of these population cycles is likely an effect of the mixotrophic diet and reduced en-ergy reserves of the individual hosts [36]. In general, when the free-living bacteria become a limiting resource, the symbionts provide a stable alternative nutritional source for the host [14, 17, 18], which does not need to rely on an energy reserve. As a consequence, when the resource is abundant, hosts do not need to build up a large energy reserve. Instead, the individual hosts can allocate more energy to reproduction. This effect is evident in the constant environment when the resource is not limiting, where symbiotic thyasirids reach higher abundances than the asymbiotic population (Fig. 2A).

Benefits of symbiosis at low resource concentrations have been documented before, theoretically and experimentally, in photo-mixotrophic and aposymbiotic organisms (e.g. in ciliates and in hydra [56, 57, 58, 59]). The literature agrees that, at low resource concentrations, mixotrophic populations have significantly higher growth rates than aposymbiotic populations, which is sufficient to prevent extinction. At high resource availability, theory suggests that the benefits of sym-biosis are in reducing loss rates [59], which agrees with our previous finding of lower somatic maintenance costs for the symbiotic *T*. cf. *gouldi* when compared to the asymbiotic *Parathyasira* [36]. The role of symbionts in building-up and performing the primary energy and carbon storage for a host has only recently been described (in the chemosymbiotic flatworm *Paracatenula* [60]). Therefore, evidence agrees with our analysis and suggests a potential role of symbiosis in mitigating environmental fluctuations in populations of mixotrophic individuals.

### 4.3. Model assumptions and limitations

Our results are related to our assumptions regarding the fecundity, mortality, and competition of the individuals in the populations. For the thyasirids of Bonne Bay, we do not have enough evidence to suggest that reproduction occurs in discrete events or that there is competition for resources between the species. There-fore, in our model, reproduction occurs continuously, and each species can graze independently of the other on the free-living bacterial resource. Likewise, for simplicity and lack of detailed information, we assumed that mortality was equal for both species in all the stages. The consequences of our assumptions could influence the population structure in our results. For example, in a population of *T. gouldi*, it has been proposed that juveniles do not uniformly predominate because early juveniles are likely to suffer a greater mortality rate, compared to late juveniles and adults [55]. Nonetheless, these simplifying assumptions are unlikely to affect the overarching pattern regarding the effect of the symbiotic strategy.

The yearly cycles experienced by both populations are a direct result of the fluctuations in temperature and its effect on the resource. We modelled the temperature oscillations according to the natural variation pattern observed in Bonne Bay. The free-living bacteria are also known to be subject to seasonal variations [25]; therefore, we used a framework derived from the Metabolic Theory of Ecology to couple the environmental temperatures to the resource abundance. Even though the abundance and digestion of the bacterial symbionts in *T*. cf. *gouldi* also show a cyclical trend [25], in our formulation, we treat the symbiont abundance as constant, considering that they provide a more stable source of nutrition. Despite our assumptions, the strength of our physiologically structured model is illustrated by the differences that we revealed between the two species’ populations.

### 4.4. Reliance on symbionts

Differences in individual energy allocation have suggested that for *T*. cf. *gouldi* the symbionts may buffer resource fluctuations [36]. Our simulations suggest that symbiotic and asymbiotic thyasirids have different adaptations to persist during winter conditions when the temperatures are low, and free-living bacteria are rare. Asymbiotic thyasirids have physiological adaptations that allow them to build a larger energy reserve, which can be mobilized more when the resource is scarce. Symbiotic thyasirids have adapted via their symbionts: the reliance on a constant supply of symbiotic bacteria buffers against the effects of seasonal lows in the abundance of free-living bacteria. Thus, for the thyasirids from Bonne Bay, both the generalist mixotrophic and the specialist diet are equally successful strategies. Our results motivate the question of why symbiotic thyasirids do not rely more on their symbionts, or equivalently, why asymbiotic thyasirids do not have a broader diet. Both alternatives could lead to individuals less sensitive to environmental fluctuations and more stable populations sizes (Fig. 4A, B). Such questions are associated with the phenotypic traits that determine resource acquisition, which are thought to be defined by an intraspecific correlation between the individual morphology, physiology and behavior. In general, it is understood that there are costs that prevent the evolution of niche generalism, for example, phylogenetic constrains [9, 13]. For symbiotic thyasirids, the gill size of the host imposes a limit to the space available for colonization by symbionts [61]. If the surface area of thyasirid gills shows a positive allometric relationship to body size, as observed in other chemosymbiotic bivalves with similarly filibranchiate gills [62], the small body size of thyasirids may prevent them from harbouring the number of symbionts that would be necessary for a greater reliance (i.e., equivalent to the symbiont dependencies observed in larger bivalves). Another likely explanation is that the costs of maintaining symbionts may rise during the winter, due to an increment in the bioirrigation necessary to control the symbiont population size or to an increase in the digestion of symbionts [29]. For asymbiotic thyasirids, there may be similar phylogenetic limitations that have prevented a change in diet and have instead promoted faster somatic growth and maturation rates. Moreover, an environment with spatial variation, such as the infaunal habitat of the thyasirids, could equally favour diet specialization [10]. The limited evidence available for the thyasirids hinders a more robust inference; however, it is clear that both strategies are successfully maintained in the community.

The ubiquity of symbiosis makes it a crucial factor that can determine the outcome of ecological and evolutionary processes [2]. Nevertheless, how symbiosis affects ecology and evolution remains mostly unknown. In this study, we show how trophic symbiosis can mitigate the effect of a seasonal environment in a population of bivalve hosts. Although we parameterized our model for the particular system of thyasirid bivalves from the fjord of Bonne Bay, Canada, our approach has a general nature, and our results are relevant in a variety of trophic symbiosis. Our results highlight the relevance of linking individual energetics and life history to population dynamics and are the first step towards a general understanding of the role of symbiosis in populations’ resilience.

## Acknowledgments

We thank the members of the Theoretical Biology Laboratory at MUN for their constructive comments on the manuscript. JM was supported by the School of Graduate Studies Baseline Fellowship from Memorial University. SCD and AH received funding from the Natural Science and Engineering Research Council of Canada (NSERC Discovery Grants 2015-06548 and 2014-05413). We also acknowledge the insightful comments from three anonymous reviewers, which helped us improve the final version of this manuscript.

## Supplementary

### Individual model

### Metabolic acceleration: DEB-abj model

In the DEB theory framework, species with larval development typically exhibit a slow embryonic development combined with a faster development during the late juvenile and adult stages [46]. This permanent increase in the metabolic rate is called ℳ metabolic acceleration. The most common type of acceleration, called acceleration, involves a simultaneous increase in the assimilation and reserve mobilization rates between birth and metamorphosis, as well as an increase in growth, maturation, reproduction and respiration. We quantified the metabolic increase in type ℳ acceleration according to the shape correction function:

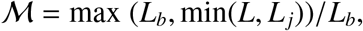

where *L* is the length in structure, and *L*_*b*_ and *L*_*j*_ represent the structural lengths at the beginning and at the end of the acceleration, which correspond to the structural lengths at birth and at metamorphosis.

### Observables: fecundity

The state variables of the DEB model are quantities that are not directly measurable. Hence, to calculate how the cumulative energy invested into reproduction (*E*_*R*_) translates to number of embryos per time, we used the following relation (Eq. 2.56 in [44]):

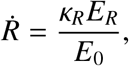

where κ_*R*_ is the reproduction efficiency (set to the standard value of 0.95) and *E*_0_ (J) is the amount of initial energy reserve invested into each embryo (i.e. the cost of an egg). To calculate *E*_0_ (Eq. 2.42 in [44]) we used the routine initial scaled reserve in the DEBtool MATLAB package [63]. The equation for 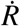 is used before calculating the cumulative reproduction of the population (equation 11).

### Grid resolution and computational time

We ran the models for Section 3.1 using a grid resolution of 30^3^ cells for ten years, measuring each day’s population density. In Section 3.2, we used a grid resolution of 20^3^ cells for six years, with output also given every day. Similarly, for Section 3.3, we integrated the model for six years with output each day, but using a grid of 15^3^ cells.

We used an Intel Xeon E5-1650 v2 @ 3.50GHz processor to run the models. We measured the integration time in core-years (calculated as hours (nodes cores) / (365 · 24)). The integration time for the models for *Thyasira* cf. *gouldi* and *Parathyasira* sp. in a constant environment at a resolution of 30^3^ cells was 0.002 and 0.012 core-years respectively. For the same grid size in the seasonal scenarios, the running time was 0.218 core-years for *T*. cf. *gouldi* and 0.232 core-years for *Parathyasira*. The simulations for *T*. cf. *gouldi* at a 20^3^ resolution took 0.018 core-years to complete, and 0.004 core-years using a grid size of 15^3^ cells.

